# Benefits and limits of biological nitrification inhibitors for plant N uptake and the environment

**DOI:** 10.1101/2023.06.07.543901

**Authors:** Christian W. Kuppe, Johannes A. Postma

**Affiliations:** Forschungszentrum Jülich GmbH, Institute of Bio- and Geosciences – Plant Sciences (IBG-2), 52425 Jülich, Germany; RWTH Aachen University, Faculty 1, Germany

**Keywords:** Rhizosphere model, bacteria, NUE, N leaching, BNI exudation

## Abstract

Does root-exudation of biological nitrification inhibitors (BNIs) facilitate nitrogen (N) uptake and reduce pollution by N loss to the environment? We modeled the spatial-temporal dynamics of nitrifiers, ammonium, nitrate, and BNIs around a root and simulated root N uptake and net rhizosphere N loss over the plant’s life cycle. We determined the sensitivity of N uptake and loss to variation in the parameters, testing a broad range of soil-plant-microbial conditions.

An increase in BNI exudation reduces net N loss and, under most conditions, plant N uptake. BNIs decrease uptake in the case of (1) low ammonium concentrations, (2) high ammonium adsorption to the soil, (3) rapid nitrate-or slow ammonium uptake by the plant, and (4) a slowly growing or fast-declining nitrifier population.

Bactericidal inhibitors facilitate uptake more than bacteriostatic ones. Some nitrification, however, is necessary to maximize uptake by both ammonium and nitrate transporter systems. Selection for increased BNI exudation should co-select for improved ammonium uptake. BNIs can reduce N uptake, which may explain why not all species exude BNIs but have a generally positive effect on the environment by increasing rhizosphere N retention.

## Introduction

The macronutrient nitrogen (N) is essential for plant metabolism amounting to 1–5 % N of the total plant dry weight ^1^. Greater N uptake promotes photosynthetic carbon fixation and plant growth. Plants take up nitrogen primarily as ammonium 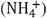 or nitrate 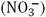 from the soil solution. At the expense of energy, plants must reduce nitrate before nitrogen can be used to synthesize amino acids. Ammonium is energetically cheaper. Nitrifying microorganisms, living in the influence sphere of the plant roots (rhizosphere), do the opposite and gain energy by converting 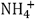 to 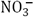 (nitrification). Nitrate is at risk of becoming unavailable to the plant and polluting the environment by leaching or nitrogen oxide emissions during denitrification. Farmers fertilize with as much as three times the amount of N in the crop yield, thus for every N in cereal crops two N are lost to the environment ^2^. Ammonium binds promptly to the soil exchange complex. Thus, it has been proposed that N is better retained in the soil in the form of ammonium, and that prolonged retention might be achieved by inhibiting nitrification ^3–5^. Synthetic nitrification inhibitors are commercially available and can be added to fertilizers, but they can potentially harm the environment and humans and have variable effectiveness ^6^. They inhibit mostly ammonia-oxidizing bacteria and have less influence on ammonia-oxidizing archaea, the dominant nitrifiers in most soils ^7^. Several plant species can exude biological nitrification inhibitors (BNIs) from their roots ^8–12^. For example, root exudates of *Sorghum bicolor* and *Brachiaria humidicola* contain BNIs ^13,14^. Recently, BNIs have been found in exudates of major cereals, rice, maize, and wheat ^12,15–17^. Exudation of BNIs has been suggested as a local and organic alternative for inhibiting nitrification and improving N uptake and reducing environmental N pollution ^18^. These BNIs are chemically diverse, including phenylpropanoids, quinones, benzoxazinoids, terpenes, and fatty alcohols ^8,16,19^. In rice and wheat, genetic variation for BNI and nitrification promotion has been reported ^12,15^. These results question the selection pressure for BNI exudation and the universal benefit of BNIs to plant N uptake. A more quantitative and mechanistic understanding of BNI functioning is necessary to understand their ecological and agronomic importance.

We ask whether nitrification inhibitors always promote N retention and N uptake or whether these effects are evident only under specific rhizosphere conditions. We answer our question through sensitivity analysis of a mechanistic spatiotemporal model of the rhizosphere (see graphical abstract).

## Material and Methods

We provide the model summary and main equations for the spatio-temporal changes here and refer to the Supplemental Information for the detailed description and parameterization. Symbols are in Table 1.

**Table 1.**
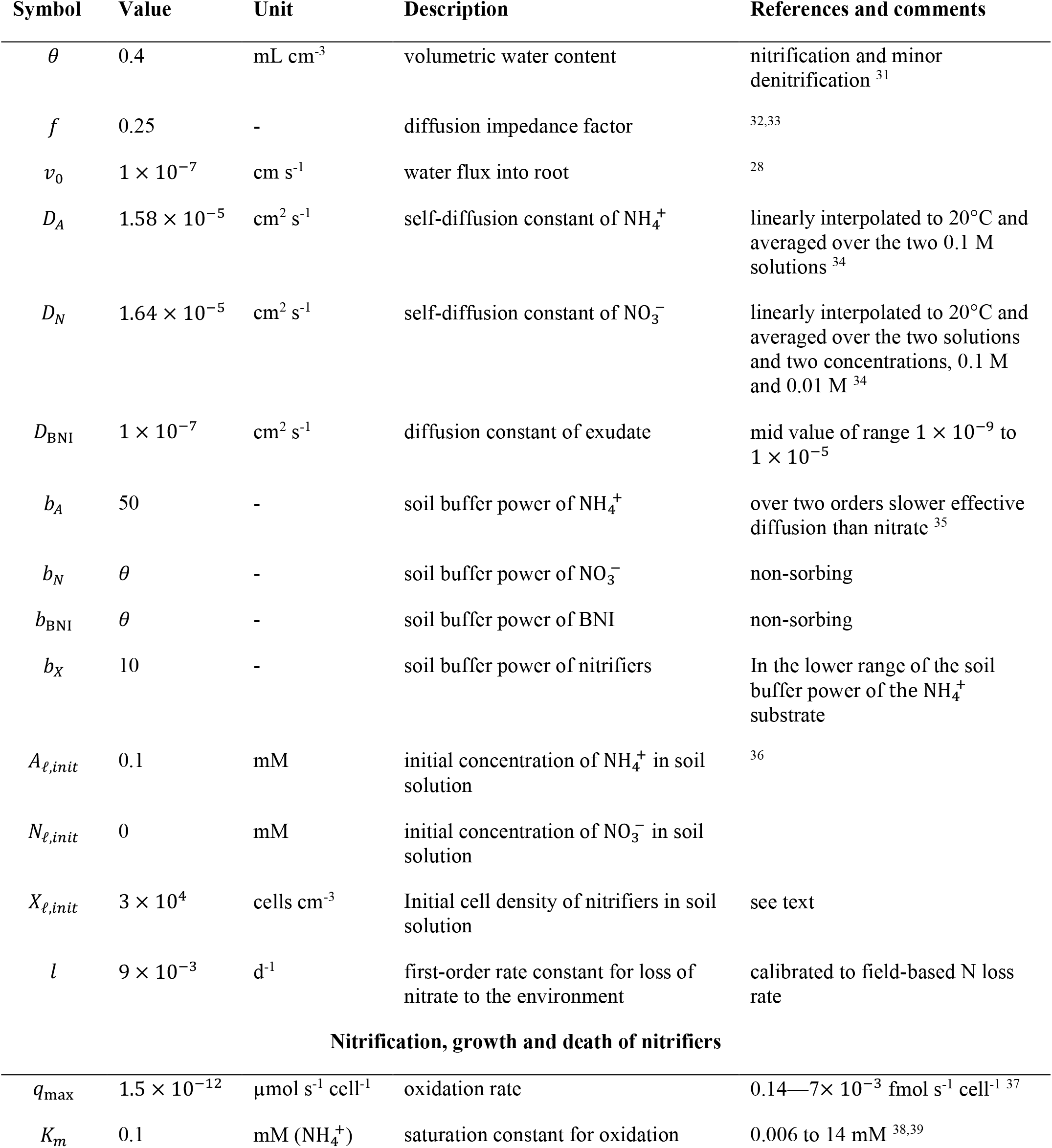
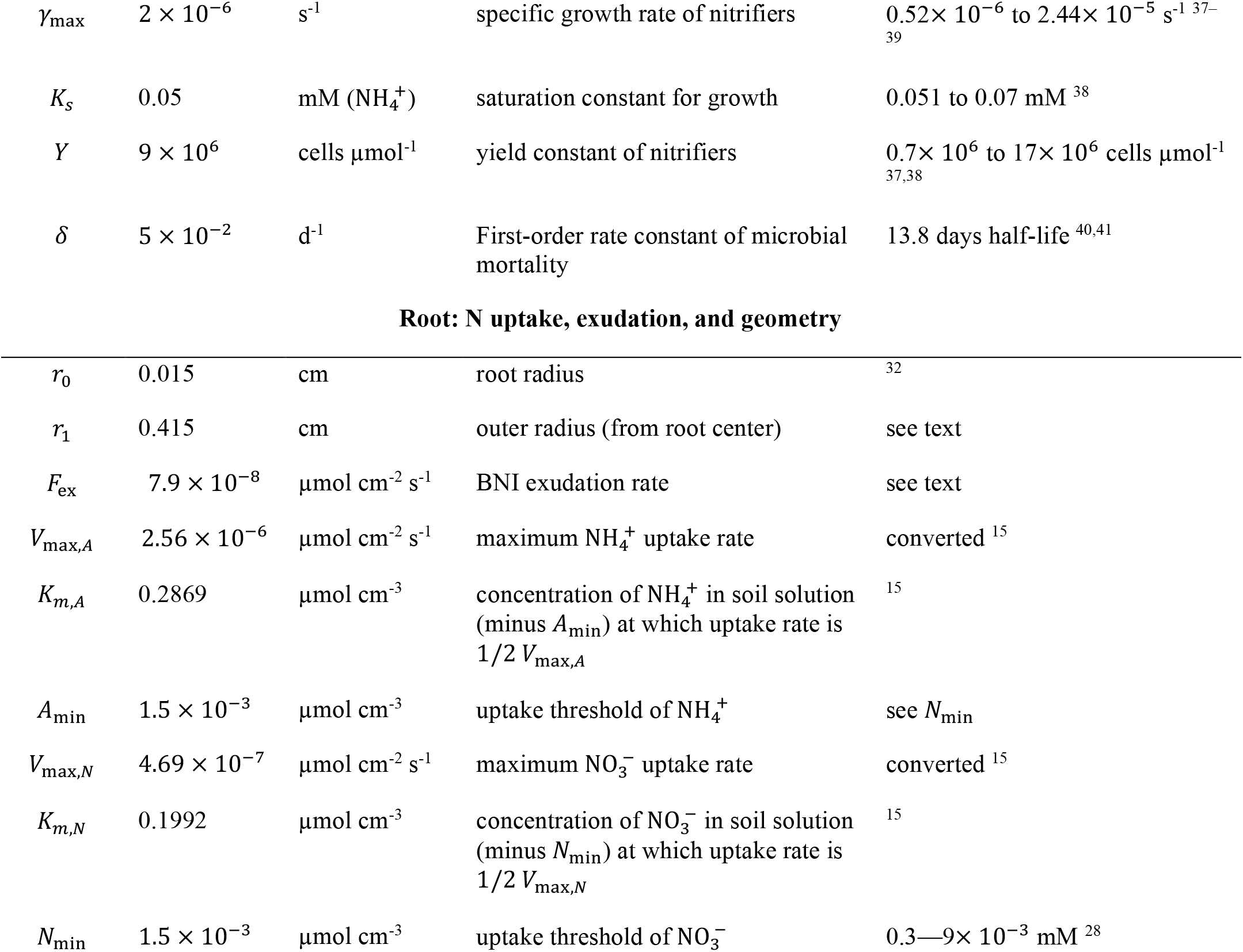
Parameters of reference simulation, their description, and literature reference

| Symbol | Value | Unit | Description | References and comments |
| --- | --- | --- | --- | --- |
| $\theta$ | 0.4 | $\text{mL cm}^{-3}$ | volumetric water content | nitrification and minor denitrification <sup>31</sup> |
| $f$ | 0.25 | - | diffusion impedance factor | 32,33 |
| $v_0$ | $1 \times 10^{-7}$ | $\text{cm s}^{-1}$ | water flux into root | 28 |
| $D_A$ | $1.58 \times 10^{-5}$ | $\text{cm}^2 \text{s}^{-1}$ | self-diffusion constant of $\text{NH}_4^+$ | linearly interpolated to 20°C and averaged over the two 0.1 M solutions <sup>34</sup> |
| $D_N$ | $1.64 \times 10^{-5}$ | $\text{cm}^2 \text{s}^{-1}$ | self-diffusion constant of $\text{NO}_3^-$ | linearly interpolated to 20°C and averaged over the two solutions and two concentrations, 0.1 M and 0.01 M <sup>34</sup> |
| $D_{\text{BNI}}$ | $1 \times 10^{-7}$ | $\text{cm}^2 \text{s}^{-1}$ | diffusion constant of exudate | mid value of range $1 \times 10^{-9}$ to $1 \times 10^{-5}$ |
| $b_A$ | 50 | - | soil buffer power of $\text{NH}_4^+$ | over two orders slower effective diffusion than nitrate <sup>35</sup> |
| $b_N$ | $\theta$ | - | soil buffer power of $\text{NO}_3^-$ | non-sorbing |
| $b_{\text{BNI}}$ | $\theta$ | - | soil buffer power of BNI | non-sorbing |
| $b_X$ | 10 | - | soil buffer power of nitrifiers | In the lower range of the soil buffer power of the $\text{NH}_4^+$ substrate |
| $A_{\ell, \text{init}}$ | 0.1 | mM | initial concentration of $\text{NH}_4^+$ in soil solution | 36 |
| $N_{\ell, \text{init}}$ | 0 | mM | initial concentration of $\text{NO}_3^-$ in soil solution | |
| $X_{\ell, \text{init}}$ | $3 \times 10^4$ | $\text{cells cm}^{-3}$ | Initial cell density of nitrifiers in soil solution | see text |
| $l$ | $9 \times 10^{-3}$ | $\text{d}^{-1}$ | first-order rate constant for loss of nitrate to the environment | calibrated to field-based N loss rate |
| **Nitrification, growth and death of nitrifiers** |  |  |  |  |
| $q_{\text{max}}$ | $1.5 \times 10^{-12}$ | $\mu\text{mol s}^{-1} \text{cell}^{-1}$ | oxidation rate | $0.14\text{--}7 \times 10^{-3} \text{ fmol s}^{-1} \text{cell}^{-1}$ <sup>37</sup> |
| $K_m$ | 0.1 | mM ($\text{NH}_4^+$) | saturation constant for oxidation | 0.006 to 14 mM <sup>38,39</sup> |
| $\gamma_{\max}$ | $2 \times 10^{-6}$ | $\text{s}^{-1}$ | specific growth rate of nitrifiers | $0.52 \times 10^{-6}$ to $2.44 \times 10^{-5} \text{ s}^{-1}$ <sup>37–39</sup> |
| $K_S$ | 0.05 | $\text{mM (NH}_4^+)$ | saturation constant for growth | 0.051 to 0.07 $\text{mM}$ <sup>38</sup> |
| $Y$ | $9 \times 10^6$ | $\text{cells } \mu\text{mol}^{-1}$ | yield constant of nitrifiers | $0.7 \times 10^6$ to $17 \times 10^6 \text{ cells } \mu\text{mol}^{-1}$ <sup>37,38</sup> |
| $\delta$ | $5 \times 10^{-2}$ | $\text{d}^{-1}$ | First-order rate constant of microbial mortality | 13.8 days half-life <sup>40,41</sup> |
| **Root: N uptake, exudation, and geometry** |  |  |  |  |
| $r_0$ | 0.015 | cm | root radius | <sup>32</sup> |
| $r_1$ | 0.415 | cm | outer radius (from root center) | see text |
| $F_{\text{ex}}$ | $7.9 \times 10^{-8}$ | $\mu\text{mol cm}^{-2} \text{ s}^{-1}$ | BNI exudation rate | see text |
| $V_{\max,A}$ | $2.56 \times 10^{-6}$ | $\mu\text{mol cm}^{-2} \text{ s}^{-1}$ | maximum $\text{NH}_4^+$ uptake rate | converted <sup>15</sup> |
| $K_{m,A}$ | 0.2869 | $\mu\text{mol cm}^{-3}$ | concentration of $\text{NH}_4^+$ in soil solution (minus $A_{\min}$) at which uptake rate is $1/2 V_{\max,A}$ | <sup>15</sup> |
| $A_{\min}$ | $1.5 \times 10^{-3}$ | $\mu\text{mol cm}^{-3}$ | uptake threshold of $\text{NH}_4^+$ | see $N_{\min}$ |
| $V_{\max,N}$ | $4.69 \times 10^{-7}$ | $\mu\text{mol cm}^{-2} \text{ s}^{-1}$ | maximum $\text{NO}_3^-$ uptake rate | converted <sup>15</sup> |
| $K_{m,N}$ | 0.1992 | $\mu\text{mol cm}^{-3}$ | concentration of $\text{NO}_3^-$ in soil solution (minus $N_{\min}$) at which uptake rate is $1/2 V_{\max,N}$ | <sup>15</sup> |
| $N_{\min}$ | $1.5 \times 10^{-3}$ | $\mu\text{mol cm}^{-3}$ | uptake threshold of $\text{NO}_3^-$ | $0.3\text{—}9 \times 10^{-3} \text{ mM}$ <sup>28</sup> |

We pose that the rooted soil domain can be represented by a root of unit length with an average associated soil volume (rhizosphere) given by the root length density (root length by soil volume) ^20^. We model the spatiotemporal dynamics of ammonium, nitrate, BNIs, and nitrifiers in this rhizosphere. Nitrifiers are modeled as one functional group oxidizing ammonium to nitrate. Root-released BNIs can reduce both the oxidation rate of ammonium to nitrate and the growth rates of the nitrifier population. Roots can take up both ammonium and nitrate, whereas the nitrifiers incorporate only ammonium into their biomass. Ammonium is the representative state variable since ammonia and ammonium are assumed to be in equilibrium. Ammonium and nitrate uptake by the root is modeled with Michaelis-Menten kinetics ^15^. Uptake of ammonium and oxidation to nitrate by nitrifiers are also saturating functions. The nitrate loss rate from the rhizosphere depends linearly on the nitrate concentration and represents the net loss of the whole soil domain, without consideration of heterogeneity at a larger scale. We simulate over a 150-day season.

Plant-N uptake can be expressed relative to the amount of N in the soil and then is typically named nitrogen use efficiency (NUE) or N recovery. We define an average rhizosphere NUE for the rooted zone in a field as

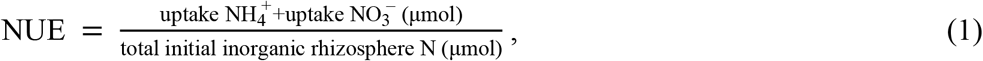

We define a relative N loss to the environment (RNL) per initial amount of N in the rhizosphere as

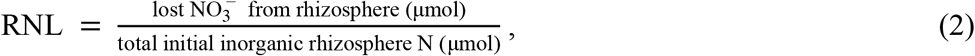

which is an average net N loss from the rooted zone in a field. We used NUE and RNL to evaluate the effect of BNIs on plant N uptake and N loss to the environment.

The changes in ammonium and nitrate concentrations in rhizosphere soil are described as

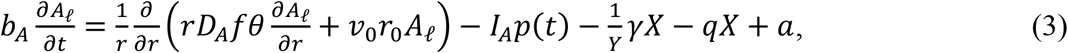

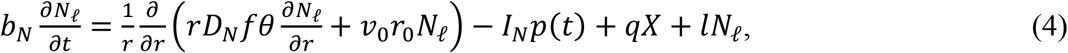

where *A*_*ℓ*_ and *N*_*ℓ*_ are ammonium and nitrate concentrations in soil solution and *b*_*A*_, *b*_*N*_ are the soil buffer powers. The nitrifier population density in soil is *X*, equation 12. The uptake of ammonium and nitrate by the root is modeled via sink terms in equations 3 and 4 for the root hairs, *I*_*A*_, *I*_*N*_, and flux-boundary conditions at the root surface. Michaelis-Menten kinetics describe the uptake of ammonium and nitrate by the root for all types of transporters. For nitrate, the flux into the root at *r* = *r*_0_ is described by the inner-boundary condition

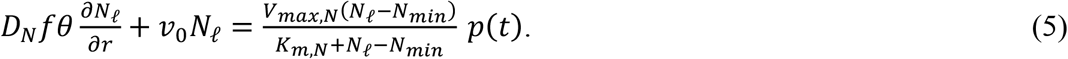

And the flux at the mid-distance to a mirrored neighboring root segment is (zero-flux outer-boundary condition at *r* = *r*_1_)

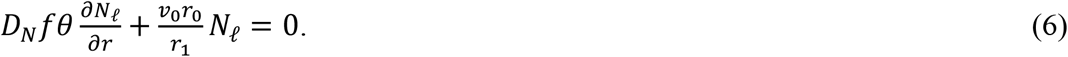

The boundary equations for ammonium are equivalent. The initial concentrations in soil solution are constant, *A*_*ℓ*,init_ and *N*_*ℓ*,init_, at *t* = 0. The arrival time of the root (*t*_*R*_ = 14) is described by switching *p*(*t*) = 0 to 1 when *t* ≥ *t*_*R*_. Nitrogen loss to the environment is *lN*_*ℓ*_. The net ammonification rate is *a* = 0 for simplification.

The relative nitrification rate is

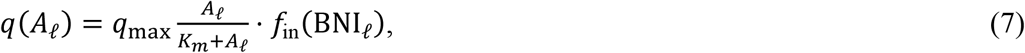

where the saturation constant for oxidation is *K*_*m*_, and maximum oxidation rate constant is *q*_max_.

The fitted nitrification inhibition function is 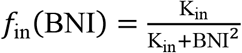 (Figure 1) and the change of BNIs is describe by

**Figure 1.**
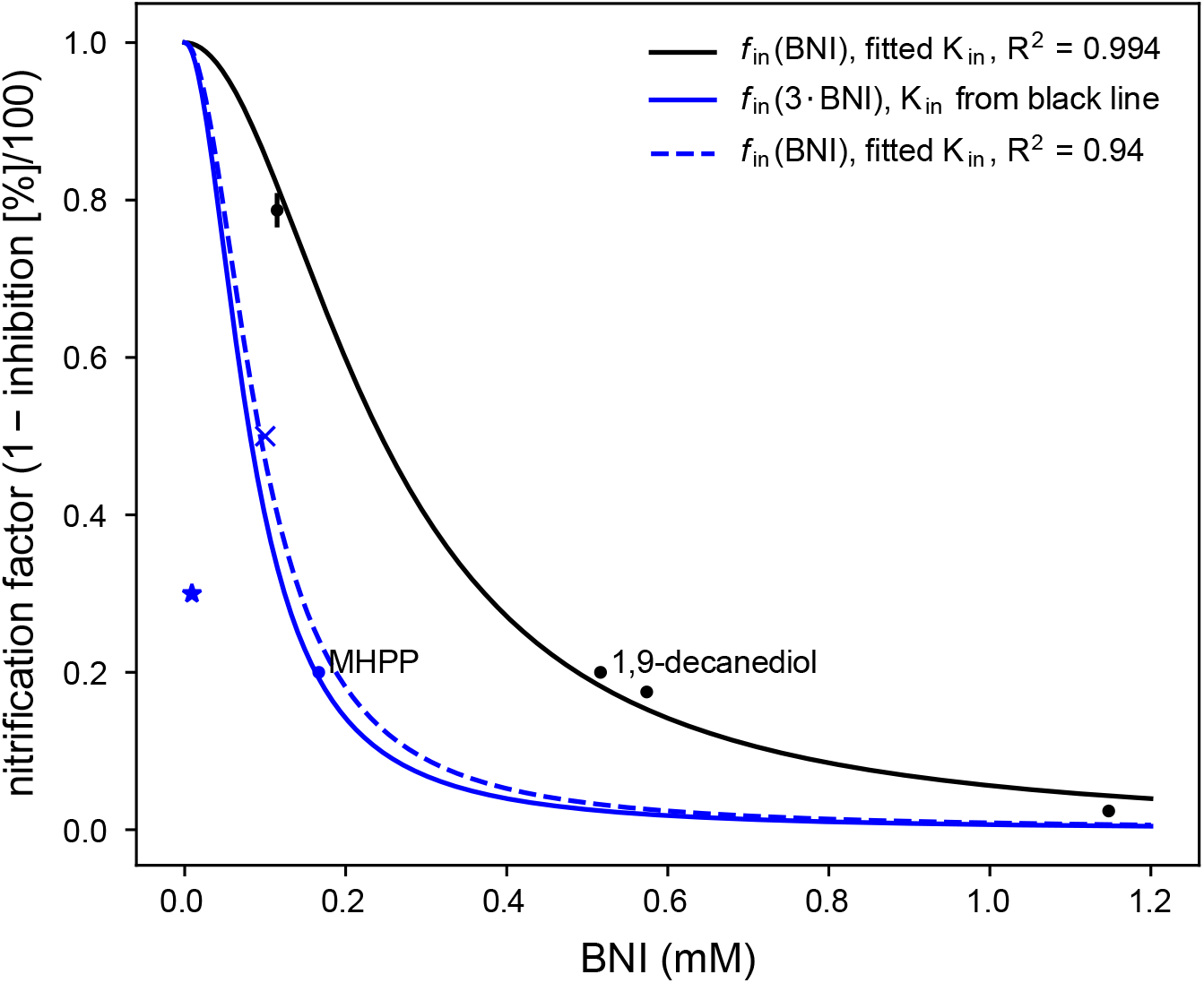
Nitrification inhibition, exemplified by the isolated BNI compounds 1,9-decanediol, black, and methyl 3-(4-hydroxyphenyl) propionate (MHPP), blue; from data by Sun et al. ^15^, ‘dots’, Subbarao et al. ^3^, ‘x’, and Zakir et al. ^13^, ‘star’. Fitting function: *f*in(BNI) = K_in_/(K_in_ + BNI^2^). The inhibition by isolated MHPP is about 2.5 to 3 times stronger than 1,9-decanediol in assay. We used K_in_ = 0.00884 (dashed line); the K_in_ for the other lines was 0.0595 mM^2^.

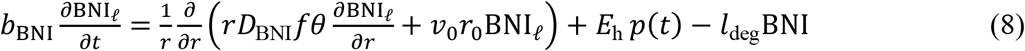

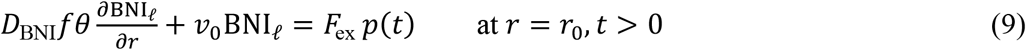

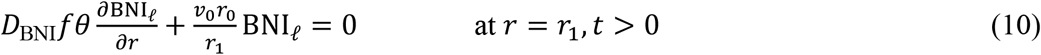

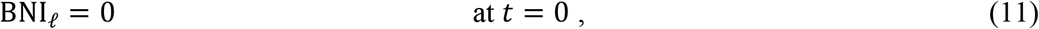

where *E*_h_ is the volumetric root hair surface area. We set *l*_deg_ = 0. The nitrifier population dynamic is modeled as ODE for each discretization point (*i*) for above’s PDE system:

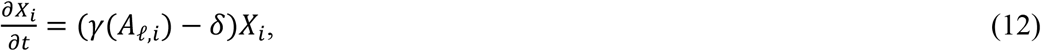

and the relative growth rate of the microbial population is described by

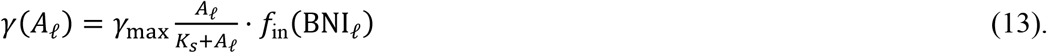

## Results

First, we established a reference simulation parameterized with values in ranges found in the literature (Table 1, Figure 2). With these reference values, the model simulated that BNIs increased total N uptake by the root segment by almost 22 %, raising the rhizosphere N use efficiency (equation 1, NUE) from 33 % to 40 % (Figure 3 blue lines and Figure 4 for multiplier =1). The relative N loss (equation 2, RNL) decreased from 49 % to 26 %, and the inorganic N converted to microbial biomass was reduced by over 43 %.

**Figure 2.**
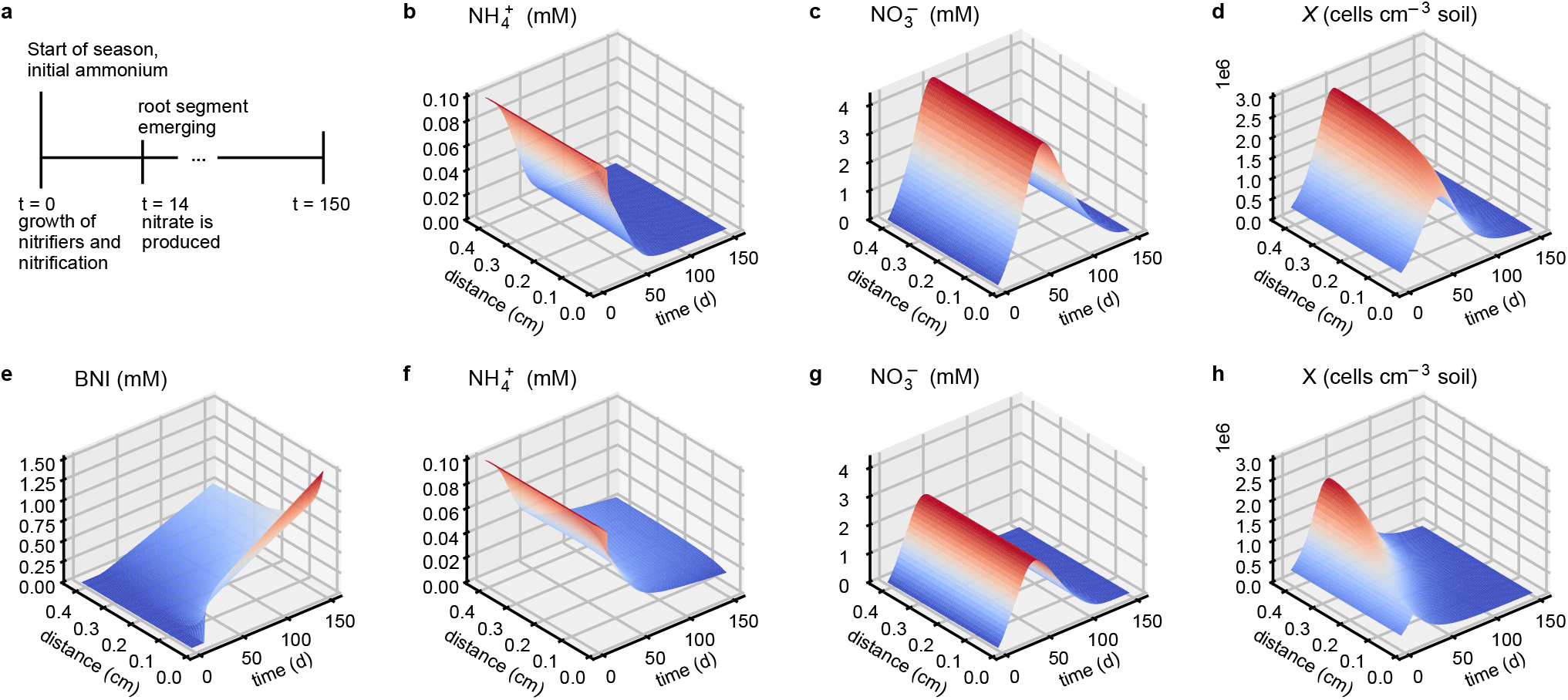
Simulation results of the reference simulation (parameter values in Table 1) without (top) and with (bottom) the exudation of BNIs. Graphs show the spatio-temporal dynamics in ammonium (b, f), nitrate (c, g), and BNI (e) concentrations in soil solution and nitrifier cell densities in soil (d, f). The distance is radial to a representative root segment of unit length (root center at distance r = 0) and the colors match the contour lines. (a) Timeline of simulated season (150 days): nitrification starts at t = 0, uptake, and exudation by the root segment at t = 14 d. Uniform growth of nitrifiers for t<14 d.

**Figure 3.**
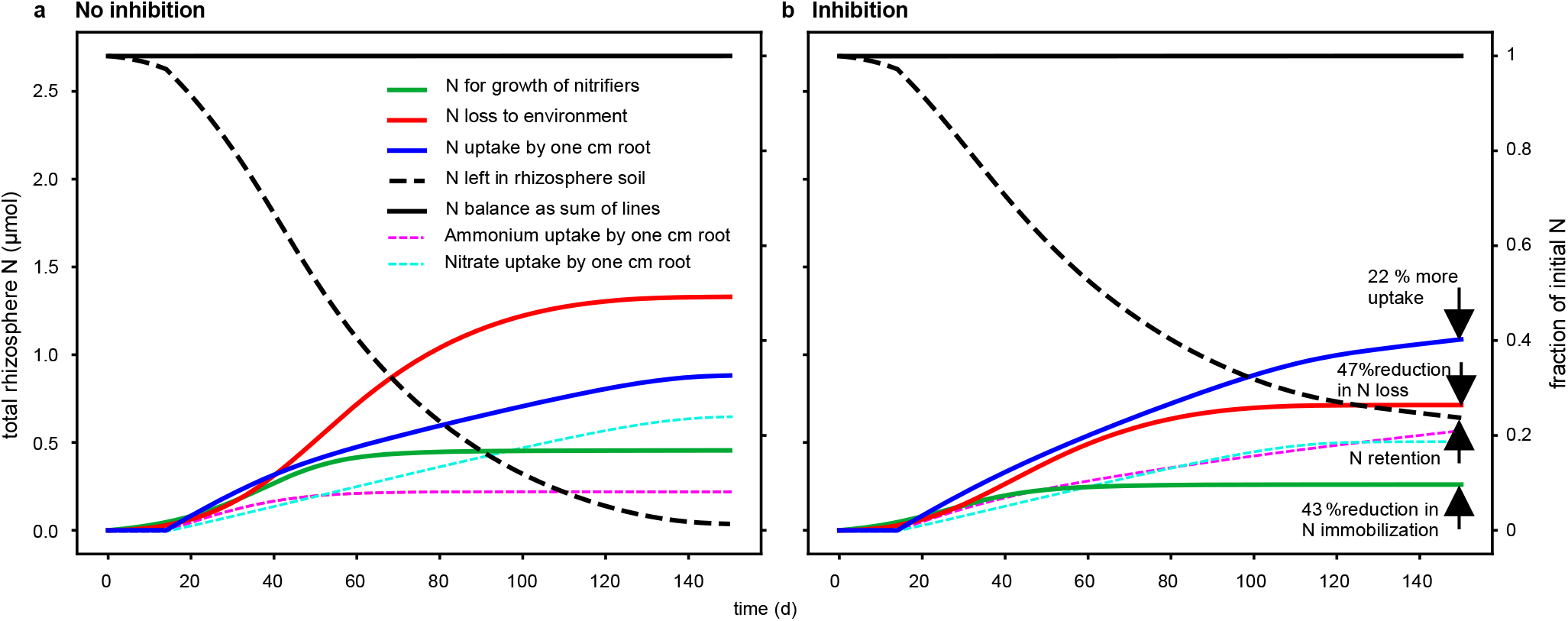
Time dynamics in the size of N contribution for simulations without (a) and with (b) inhibition. When BNIs are exuded, N uptake is increased, and N loss and immobilization are reduced. At the end, there is nitrogen left in the rhizosphere, suggesting fertilization can be reduced. The dashed lines show inorganic N retained in the rhizosphere due to adsorption. Parameter values are in Table 1. Distinction between uptake by hairs and main root surface is shown in supplemental Figure S2.

**Figure 4.**
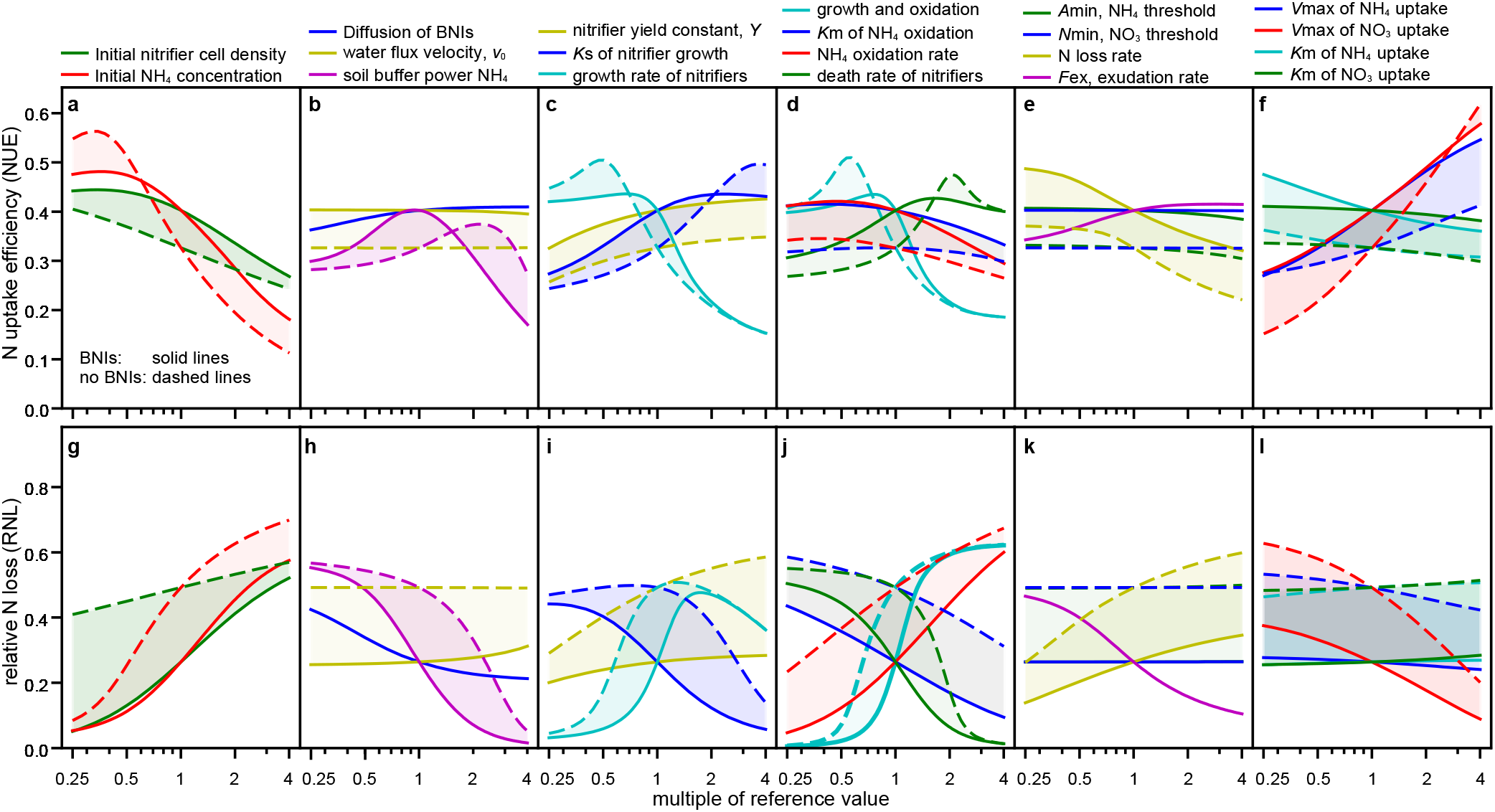
Sensitivity of N uptake efficiency (first row) and relative loss (second row) to variation in parameter values. The shading indicates the difference between simulations with inhibition by exudation (solid lines), and without (dashed lines). The x-axes indicate the change in the parameter value relative to the value of the reference simulation. Note that the x-axis is logarithmic and a 1:1-line is not straight in a semi-log-plot. The relative change in sensitivity (steepness of curves) can be compared between all sub-plots since the y-axes of each sub-plot have the same relative scaling to their reference simulation without inhibition. Note that the initial N concentration varies only in (a) and (g). BNIs increase uptake, except if NO ^−^ concentration is low, or highly sorbing, 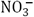 uptake rate is high or 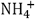 uptake rate is low, nitrifiers grow slowly, or their death rate is high, or both nitrifier growth and oxidation are slow. However, if the death rate is too high, the reduction of nitrifier population is too fast such that nitrifier and their inhibition does not matter (d, j). BNIs always reduce RNL (g—l).

Next, we varied soil, microbial, and plant conditions to show when BNI exuding plants have an advantage. We performed a sensitivity analysis on NUE and RNL over a 16-fold change in parameter values (Figure 4). In Figure 4, the inhibition gap is the difference between the results without (dashed lines) and with inhibition (solid lines). The difference between N uptake with and without inhibition can swap signs, indicating inhibition affected uptake negatively. In all scenarios, BNIs reduced RNL, although this benefit of BNIs varied strongly. NUE was often improved by BNIs, but not always. We summarize the results by looking at the governing processes that influence ammonium and nitrate uptake by the root.

### BNIs are beneficial when ammonium concentrations are large

The initial ammonium concentration determines whether BNIs increase or decrease NUE (Figure 4a, red lines, also supplemental Figure S1). For example, the same N uptake as the reference simulation (without inhibition) can be reached with 25% less initial ammonium when the root exudes BNIs and nitrate loss over the season would be reduced by 70% (Figure S1). Low initial ammonium concentration increases NUE in all conditions. The rhizosphere ammonium concentration is not depleted, with and without BNIs. However, inhibition of nitrate production by BNIs decreased total N uptake and NUE. A greater uptake was achieved when both nitrate and ammonium were available in the rhizosphere, such that both uptake transporter systems take up N at elevated rates (Pareto efficiency). Low initial ammonium decreases loss because nitrification is slow, and the plant takes up the produced nitrate before it is lost. In these scenarios, further inhibition of nitrification by BNI, on top of the intrinsic inhibition caused by the low initial ammonium concentrations, does not reduce loss further but reduces uptake of nitrogen. Inhibition affected N loss to a lesser extent (smaller gap) if the initial cell densities of nitrifiers were high (Figure 4g).

### BNIs are beneficial when the adsorption of ammonium to the soil is low

The change in soil buffer power ^20^ of ammonium (=dissolved/total NH_4_) includes a change in its initial concentration in the soil solution to have the same total initial concentrations. A decrease in soil buffer power decreased N uptake by the plant and increased loss from the rhizosphere (without and with inhibition, Figure 4b, h). Despite the larger initial concentration in the soil solution, ammonium was depleted faster with reduced soil buffer power. An increase in soil buffer power of ammonium decreased ammonium in solution and, consequently, nitrate production. Hence, compared with the reference simulation, this nitrate concentration in the rhizosphere was depleted faster, and the root’s nitrate uptake system became under-utilized. Nitrifiers replenish the nitrate concentration in the rhizosphere when not inhibited. Hence, in soils that strongly adsorb ammonium, BNIs are not beneficial.

Sorption reduces the effective diffusion rates and thereby the concentration of BNIs throughout the rhizosphere. Slow effective diffusion of BNIs causes BNIs to accumulate at the root surface, reducing their overall effectiveness across the rhizosphere (Figure 4b, h, blue line).

The reference values for BNI exudation rate and soil buffer power resulted in near optimal NUE (Table 1, Figure 4b, h, pink lines). Increasing the rate of BNI exudation, *F*_ex_, leads to saturation of the relative effectiveness of BNIs on uptake, which indicates a good balance in the reference simulation between metabolic cost of BNI exudation and its utility to increase N uptake. N uptake and loss were insensitive to variation in the water flux velocity, *v*_0_, caused by the transpiration stream (Figure 4h).

### BNIs are beneficial when the nitrifier population growth is neither fast nor slow

The effect of BNIs on NUE and RNL was sensitive to changes in the parameter values of nitrifier growth (*γ*_max_ and *K*_*s*_, equation 13; Figure 4c, i, cyan lines). Fast growth led to immobilization of ammonium, and the consequent larger nitrifier population oxidized the remaining ammonium rapidly. The resulting low ammonium concentrations limited nitrification, and thereby, made BNIs redundant once exuded. When the ammonium concentration in the rhizosphere depleted, roots and nitrifiers competed for ammonium. This competition reduced the total N uptake by the plant. If the nitrifier population grew too slowly, BNIs did not facilitate uptake either because nitrate production was insufficient to maintain total N uptake. Thus, both low and high nitrifier growth rates lead to BNIs being less effective.

The yield constant of nitrifiers, *Y*, relates the population size to ammonium concentrations, affecting the ammonium uptake by nitrifiers. Changes in this parameter value influenced N loss more than N uptake by the root.

### BNIs are beneficial when nitrifiers are persistent in soils

For high death rates, there was no difference with or without exudation because nitrifiers died early (Figure 4d, j, green lines) without contributing to the inorganic ammonium concentration. For variation in death and growth rates, uptake had optima. Hence, the nitrifier population should not be too low, producing sufficient nitrate for uptake by the plant over the season.

### Uptake is insensitive to changes in oxidation rates

The nitrification rate, *q*, increases with increasing maximum oxidation coefficient, *q*_max_, and decreasing saturation constant for oxidation, *K*_*m*_ (equation 7). Nitrogen uptake and loss were less sensitive to variation in nitrification rates than death rates (Figure 4d, j, compare blue and red to green lines). Biological nitrification inhibitors function on both processes, but the greater sensitivity for death rates implies that control over the nitrifier population is the more important function. Increased death rates without BNIs exceeded the effect of BNIs because controlling population by BNIs has a delay.

### BNIs decrease the sensitivity to nitrate loss rate

The sensitivity of NUE and RNL regarding changes in the first-order rate coefficient of N loss (*l*) is almost reciprocal to changes in the exudation rate. With BNI exudation, the RNL was more robust to nitrate loss. Biological nitrification inhibitors are slightly more effective for slower loss (Figure 4e, k, yellow lines).

### Uptake kinetics of the root determine the efficacy of BNIs

Nitrogen uptake and loss were insensitive to changes in the minimum concentrations for uptake of ammonium and nitrate by the root (*A*_min_, *N*_min_, equation 5) but BNI efficacy was affected by the other kinetic parameters, *V*_max_ and *K*_*m*_. When the plant is able to take up nitrate very fast (large *V*_max,*N*_), inhibition did not facilitate uptake. The opposite is true for roots with fast ammonium uptake (large *V*_max,*A*_, Figure 4f). The lines for the affinity constants *K*_*m,N*_ and *K*_*m,A*_ are less sensitive than *V*_max,*N*_ and *V*_max,*A*_. Thus, whether BNIs are beneficial for uptake depends on soil conditions as well as on the uptake transporters of the root. Taken together, these results indicate that the biggest gain in plant nitrogen uptake is expected by inhibiting growth of nitrifiers and, concurrently, by increasing plant ammonium uptake kinetics.

## Discussion

We asked if and under what conditions BNIs improve both NUE and RNL. BNI exudation benefits uptake often, but not in all scenarios. Our modeling results support previous suggestions that BNI exudation can increase NUE, especially when plants take up ammonium faster than nitrate, as is the case for many rice genotypes ^15,21^.

The time dynamics of rhizosphere ammonium and nitrate concentrations ultimately determine the efficacy of BNIs for plant N uptake and N loss. Determining processes are 1) competition for nitrogen between plant roots and microbes, 2) the rate of depletion of ammonium, and 3) the utilization of the root’s nitrate transporters (synergistic uptake ^9^). Low ammonium concentration lowers nitrification rates, and no further inhibition is necessary. At larger ammonium concentrations, nitrification increases total N uptake by increasing the nitrate concentrations if the ammonium uptake is not reduced by equal amounts. This relationship could explain why exudates of some rice varieties promote nitrification ^15^.

BNI exudation has a metabolic cost to the plant and, as we showed here, does not always increase total N uptake, which may explain why BNIs are not found in all species ^22^. In environments that are less favorable for the nitrifiers, the reduced growth and longevity makes BNIs redundant. The same is true in low nitrate and ammonium environments, which result in an intrinsic inhibition by the low concentrations. Selection pressure in these environments might have been low. However, BNI production was found in species adapted to low-N conditions ^9^. This seems to contradict the simulation results and needs further investigation. Increased initial ammonium concentrations typically occur under agricultural fertilized conditions, which makes the ability of releasing BNIs a breeding target.

In some species, ammonium uptake causes an excess uptake of cations over anions, and this imbalance triggers BNI exudation ^23–25^. However, instead of an ammonium dependent exudation, our model necessitates negative feedback to nitrate concentrations, because the root’s nitrate transport system can be under-utilized when BNIs exudation reduces nitrate production. If both ammonium and nitrate are sufficiently available to saturate uptake over the growth period, nitrification inhibition does not affect N uptake. Future experiments could test whether uptake-saturating ammonium concentrations promote, and low nitrate concentrations inhibit BNI exudation balancing uptake and investment by the plant. This feedback would also address the question if an exudate inhibits specifically ^9^.

Exudates might reduce the enzymatic oxidation or the nitrifier population size. BNIs from tree roots slow down nitrifier growth, while those from wheat reduce ammonium oxidation ^11,26^. Many experiments, however, do not distinguish death, growth, and oxidation. In bacteria assays, addition of the intermediate substrate hydroxylamine only partially rescued the nitrification process inhibited by BNIs ^15,22^. This means either that not all enzymes were inhibited or that bacteria died. Thus, the inhibition of enzymes was proposed ^3,14^, but that does not rule out other modes of action ^8^, as fatty alcohols can potentially damage the cell membranes of bacteria ^7^. Bacteria might be inactivated by the BNI and potentially killed (bactericidal). Our simulations indicate the latter would be more beneficial to the plant as it avoids N immobilization into the bacterial biomass. A bactericidal function, however, might affect other microbes as well, with unknown consequences.

Our model assumes that initial ammonium concentration is converted by nitrifiers, such that nitrate is present at root emergence. This scenario may represent a broad range of starting conditions: application of urea or ammonium fertilizer and zero nitrate pre-season, or a soil with a high mineralization rate after a cold or dry season. During the season, root length density increases such that the field can be defined as rhizosphere quickly. The sensitivity analysis, for example for the nitrifier death rate parameter, affected the simulation over the whole period, whereas BNIs, which also affect the population dynamics, only became effective later during the simulation. Thus, NUE and RNL were more sensitive to change in the parameter values than to BNIs and might indicate that synthetic nitrification inhibitors can be more effective as they work earlier. On the other hand, BNIs continue to be exuded throughout the season, whereas the synthetic ones must be degradable. This suggests that both forms could be complementary.

We kept the size of the rhizosphere constant, which implies a constant root length density. This assumption might be reasonable under ecological conditions and pastures with continuous ground cover over time. Annual crops are typically planted in fields, where root length density might vary over time strongly. Root length density typically increases with nitrate and ammonium availability, as plants grow faster ^27^. Future studies will need to determine the importance of these scenarios for the efficiency of BNIs. However, because BNIs are chemically diverse, we regard addressing current uncertainty in the diffusion coefficient and mode of action of BNIs as a higher priority. We took determined parameter values from published measurements, and the simulated rate of ammonium oxidation seem within range of reported values for nitrate production and its inhibition, still much uncertainty remains to their precise values in relation to different environmental factors such as soil temperature and water content ^7,28–30^.

In conclusion: BNI exudation should go hand in hand with enhanced ammonium uptake. Nitrifying microorganisms are competitors for ammonium, but they are necessary to produce some nitrate to maximize the use of both ammonium and nitrate uptake systems. Controlling the population size of nitrifiers affects total N uptake more than controlling their nitrification activity. This result suggests that exudation should be bactericidal and not bacteriostatic – kill nitrifiers but not too many.

An increase in BNI concentration reduces N loss but does not always increase N uptake. BNIs even decrease uptake when 1) the (initial) soil ammonium concentration is low, 2) the ammonium soil adsorption is high, 3) nitrate uptake is fast relative to ammonium uptake, or 4) the nitrifier population grows slowly, self-competes, or declines fast. Nitrogen uptake by the root is less sensitive to changes in oxidation rates of ammonium by nitrifiers.

Inhibition can only facilitate N uptake and uptake efficiency when the inferred lower nitrate production does not reduce the plant’s nitrate uptake. With inhibition, there is potentially longer sustained uptake of ammonium over time, but that cannot compensate for the reductions in uptake when the nitrate pool is depleted early. Nitrogen is a pollutant, and BNIs may avoid N loss while increasing NUE. Thereby, BNIs indirectly promote carbon fixation and crop production.

## ASSOCIATED CONTENT

### Supporting Information

PDF with Text and Figures.

### Data availability statement

Upon publication, all model results will be made available in a data repository.

## ACKNOWLEDGMENT

We thank Drs. Hendrik Poorter, Fabio Fiorani, Davide Bulgarelli, and Timothy George for helpful comments on the manuscript. We thank Prof. Dr. Björn Usadel and Dr. Marie Bolger for providing access and computational time on the IBG-4 cluster.

## SYNOPSIS

Nitrate is a pollutant and product of nitrification by soil microorganisms. Plants can inhibit nitrification, which reduces N loss to the environment, but not necessarily increases plant N uptake.

## Graphical Abstract

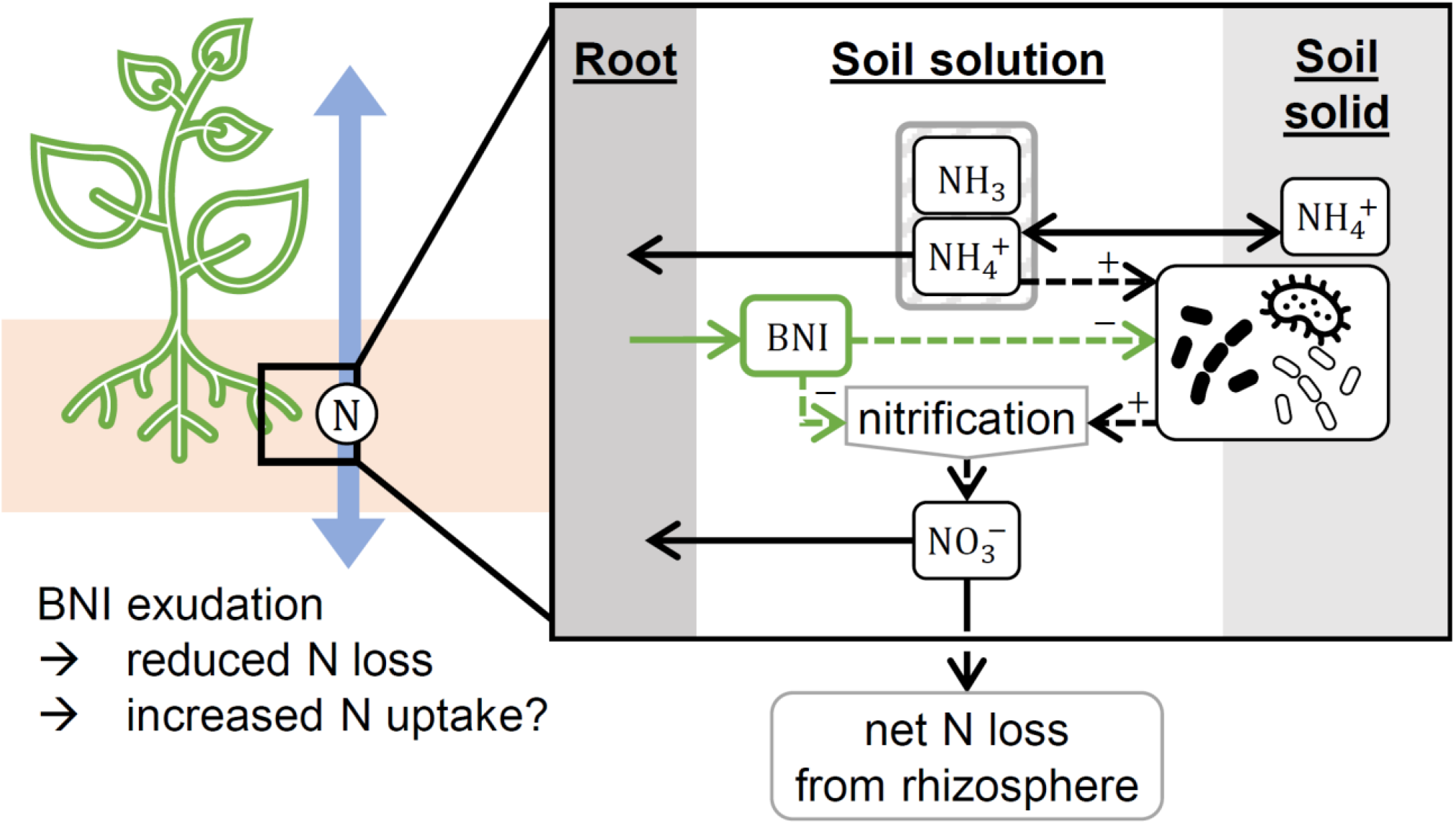

## REFERENCES

(1) Hawkesford, M.; Horst, W.; Kichey, T.; Lambers, H.; Schjoerring, J.; Møller, I. S.; White, P. Chapter 6 - Functions of Macronutrients. In Marschner’s Mineral Nutrition of Higher Plants (Third Edition); Marschner, P., Ed.; Academic Press: San Diego, 2012; pp 135–189. https://doi.org/10.1016/B978-0-12-384905-2.00006-6.

(2) Yu, X.; Keitel, C.; Zhang, Y.; Wangeci, A. N.; Dijkstra, F. A. Global Meta-Analysis of Nitrogen Fertilizer Use Efficiency in Rice, Wheat and Maize. Agriculture, Ecosystems & Environment 2022, 338, 108089. https://doi.org/10.1016/j.agee.2022.108089.

(3) Subbarao, G. V.; Nakahara, K.; Ishikawa, T.; Ono, H.; Yoshida, M.; Yoshihashi, T.; Zhu, Y.; Zakir, H. A. K. M.; Deshpande, S. P.; Hash, C. T.; Sahrawat, K. L. Biological Nitrification Inhibition (BNI) Activity in Sorghum and Its Characterization. Plant Soil 2013, 366 (1), 243–259. https://doi.org/10.1007/s11104-012-1419-9.

(4) Upadhyayl, R. K.; Patra, D. D.; Tewari, S. K. Natural Nitrification Inhibitors for Higher Nitrogen Use Efficiency, Crop Yield, and for Curtailing Global Warming. 2011, 6.

(5) Shen, J.; Li, C.; Mi, G.; Li, L.; Yuan, L.; Jiang, R.; Zhang, F. Maximizing Root/Rhizosphere Efficiency to Improve Crop Productivity and Nutrient Use Efficiency in Intensive Agriculture of China. J Exp Bot 2013, 64 (5), 1181–1192. https://doi.org/10.1093/jxb/ers342.

(6) Woodward, E. E.; Edwards, T. M.; Givens, C. E.; Kolpin, D. W.; Hladik, M. L. Widespread Use of the Nitrification Inhibitor Nitrapyrin: Assessing Benefits and Costs to Agriculture, Ecosystems, and Environmental Health. Environ. Sci. Technol. 2021, 55 (3), 1345–1353. https://doi.org/10.1021/acs.est.0c05732.

(7) Lu, Y.; Zhang, X.; Jiang, J.; Kronzucker, H. J.; Shen, W.; Shi, W. Effects of the Biological Nitrification Inhibitor 1,9-Decanediol on Nitrification and Ammonia Oxidizers in Three Agricultural Soils. Soil Biology and Biochemistry 2019, 129, 48–59. https://doi.org/10.1016/j.soilbio.2018.11.008.

(8) Nardi, P.; Laanbroek, H. J.; Nicol, G. W.; Renella, G.; Cardinale, M.; Pietramellara, G.; Weckwerth, W.; Trinchera, A.; Ghatak, A.; Nannipieri, P. Biological Nitrification Inhibition in the Rhizosphere: Determining Interactions and Impact on Microbially Mediated Processes and Potential Applications. FEMS Microbiology Reviews 2020, 44 (6), 874–908. https://doi.org/10.1093/femsre/fuaa037.

(9) Coskun, D.; Britto, D. T.; Shi, W.; Kronzucker, H. J. Nitrogen Transformations in Modern Agriculture and the Role of Biological Nitrification Inhibition. Nature Plants 2017, 3 (6), nplants201774. https://doi.org/10.1038/nplants.2017.74.

(10) Subbarao, G. V.; Kishii, M.; Bozal-Leorri, A.; Ortiz-Monasterio, I.; Gao, X.; Ibba, M. I.; Karwat, H.; Gonzalez-Moro, M. B.; Gonzalez-Murua, C.; Yoshihashi, T.; Tobita, S.; Kommerell, V.; Braun, H.-J.; Iwanaga, M. Enlisting Wild Grass Genes to Combat Nitrification in Wheat Farming: A Nature-Based Solution. Proceedings of the National Academy of Sciences 2021, 118 (35), e2106595118. https://doi.org/10.1073/pnas.2106595118.

(11) Laffite, A.; Florio, A.; Andrianarisoa, K. S.; Creuze des Chatelliers, C.; Schloter-Hai, B.; Ndaw, S. M.; Periot, C.; Schloter, M.; Zeller, B.; Poly, F.; Le Roux, X. Biological Inhibition of Soil Nitrification by Forest Tree Species Affects Nitrobacter Populations. Environmental Microbiology 2020, 22 (3), 1141–1153. https://doi.org/10.1111/1462-2920.14905.

(12) O’Sullivan, C. A.; Fillery, I. R. P.; Roper, M. M.; Richards, R. A. Identification of Several Wheat Landraces with Biological Nitrification Inhibition Capacity. Plant Soil 2016, 404 (1), 61–74. https://doi.org/10.1007/s11104-016-2822-4.

(13) Zakir, H. A. K. M.; Subbarao, G. V.; Pearse, S. J.; Gopalakrishnan, S.; Ito, O.; Ishikawa, T.; Kawano, N.; Nakahara, K.; Yoshihashi, T.; Ono, H.; Yoshida, M. Detection, Isolation and Characterization of a Root-Exuded Compound, Methyl 3-(4-Hydroxyphenyl) Propionate, Responsible for Biological Nitrification Inhibition by Sorghum (Sorghum Bicolor). New Phytologist 2008, 180 (2), 442–451. https://doi.org/10.1111/j.1469-8137.2008.02576.x.

(14) Subbarao, G. V.; Nakahara, K.; Hurtado, M. P.; Ono, H.; Moreta, D. E.; Salcedo, A. F.; Yoshihashi, A. T.; Ishikawa, T.; Ishitani, M.; Ohnishi-Kameyama, M.; Yoshida, M.; Rondon, M.; Rao, I. M.; Lascano, C. E.; Berry, W. L.; Ito, O. Evidence for Biological Nitrification Inhibition in Brachiaria Pastures. PNAS 2009, 106 (41), 17302–17307. https://doi.org/10.1073/pnas.0903694106.

(15) Sun, L.; Lu, Y.; Yu, F.; Kronzucker, H. J.; Shi, W. Biological Nitrification Inhibition by Rice Root Exudates and Its Relationship with Nitrogen-Use Efficiency. New Phytologist 2016, 212 (3), 646–656. https://doi.org/10.1111/nph.14057.

(16) Otaka, J.; Subbarao, G. V.; Ono, H.; Yoshihashi, T. Biological Nitrification Inhibition in Maize—Isolation and Identification of Hydrophobic Inhibitors from Root Exudates. Biol Fertil Soils 2021. https://doi.org/10.1007/s00374-021-01577-x.

(17) Chen, S.; He, M.; Zhao, C.; Wang, W.; Zhu, Q.; Dan, X.; He, X.; Meng, L.; Zhang, S.; Cai, Z.; Zhang, J.; Müller, C. Rice Genotype Affects Nitrification Inhibition in the Rhizosphere. Plant Soil 2022. https://doi.org/10.1007/s11104-022-05609-9.

(18) Subbarao, G. V.; Searchinger, T. D. A “More Ammonium Solution” to Mitigate Nitrogen Pollution and Boost Crop Yields. Proceedings of the National Academy of Sciences 2021, 118 (22), e2107576118. https://doi.org/10.1073/pnas.2107576118.

(19) Coskun, D.; Britto, D. T.; Shi, W.; Kronzucker, H. J. How Plant Root Exudates Shape the Nitrogen Cycle. Trends in Plant Science 2017, 22 (8), 661–673. https://doi.org/10.1016/j.tplants.2017.05.004.

(20) Kuppe, C. W.; Schnepf, A.; von Lieres, E.; Watt, M.; Postma, J. A. Rhizosphere Models: Their Concepts and Application to Plant-Soil Ecosystems. Plant Soil 2022, 474 (1), 17–55. https://doi.org/10.1007/s11104-021-05201-7.

(21) Balkos, K. D.; Britto, D. T.; Kronzucker, H. J. Optimization of Ammonium Acquisition and Metabolism by Potassium in Rice (Oryza Sativa L. Cv. IR-72). Plant, Cell & Environment 2010, 33 (1), 23–34. https://doi.org/10.1111/j.1365-3040.2009.02046.x.

(22) Subbarao, G. V.; Rondon, M.; Ito, O.; Ishikawa, T.; Rao, I. M.; Nakahara, K.; Lascano, C.; Berry, W. L. Biological Nitrification Inhibition (BNI)—Is It a Widespread Phenomenon? Plant Soil 2007, 294 (1), 5–18. https://doi.org/10.1007/s11104-006-9159-3.

(23) Subbarao, G. V.; Wang, H. Y.; Ito, O.; Nakahara, K.; Berry, W. L. NH4+ Triggers the Synthesis and Release of Biological Nitrification Inhibition Compounds in Brachiaria Humidicola Roots. Plant Soil 2007, 290 (1–2), 245–257. https://doi.org/10.1007/s11104-006-9156-6.

(24) Zhang, X.; Lu, Y.; Yang, T.; Kronzucker, H. J.; Shi, W. Factors Influencing the Release of the Biological Nitrification Inhibitor 1,9-Decanediol from Rice (Oryza Sativa L.) Roots. Plant Soil 2019, 436 (1), 253–265. https://doi.org/10.1007/s11104-019-03933-1.

(25) Egenolf, K.; Verma, S.; Schöne, J.; Klaiber, I.; Arango, J.; Cadisch, G.; Neumann, G.; Rasche, F. Rhizosphere PH and Cation-Anion Balance Determine the Exudation of Nitrification Inhibitor 3-Epi-Brachialactone Suggesting Release via Secondary Transport. Physiologia Plantarum 2021, 172 (1), 116–123. https://doi.org/10.1111/ppl.13300.

(26) Jáuregui, I.; Vega-Mas, I.; Delaplace, P.; Vanderschuren, H.; Thonar, C. An Optimized Hydroponic Pipeline for Large-Scale Identification of Wheat Genotypes with Resilient Biological Nitrification Inhibition Activity. New Phytologist 2023, n/a. https://doi.org/10.1111/nph.18807.

(27) Lopez, G.; Ahmadi, S. H.; Amelung, W.; Athmann, M.; Ewert, F.; Gaiser, T.; Gocke, M. I.; Kautz, T.; Postma, J.; Rachmilevitch, S.; Schaaf, G.; Schnepf, A.; Stoschus, A.; Watt, M.; Yu, P.; Seidel, S. J. Nutrient Deficiency Effects on Root Architecture and Root-to-Shoot Ratio in Arable Crops. Frontiers in Plant Science 2023, 13.

(28) Barber, S. A. Soil Nutrient Bioavailability: A Mechanistic Approach; John Wiley & Sons, 1995.

(29) Myrold, D. D.; Tiedje, J. M. Simultaneous Estimation of Several Nitrogen Cycle Rates Using 15N: Theory and Application. Soil Biology and Biochemistry 1986, 18 (6), 559–568. https://doi.org/10.1016/0038-0717(86)90076-3.

(30) Højberg, O.; Binnerup, S. J.; Sørensen, J. Potential Rates of Ammonium Oxidation, Nitrite Oxidation, Nitrate Reduction and Denitrification in the Young Barley Rhizosphere. Soil Biology and Biochemistry 1996, 28 (1), 47–54. https://doi.org/10.1016/0038-0717(95)00119-0.

(31) Hwang, S.; Hanaki, K. Effects of Oxygen Concentration and Moisture Content of Refuse on Nitrification, Denitrification and Nitrous Oxide Production. Bioresource Technology 2000, 71 (2), 159–165. https://doi.org/10.1016/S0960-8524(99)90068-8.

(32) Kuppe, C. W.; Kirk, G. J. D.; Wissuwa, M.; Postma, J. A. Rice Increases Phosphorus Uptake in Strongly Sorbing Soils by Intra-Root Facilitation. Plant, Cell & Environment 2022, 45 (3), 884–899. https://doi.org/10.1111/pce.14285.

(33) Barraclough, P. B.; Tinker, P. B. The Determination of Ionic Diffusion Coefficients in Field Soils. I. Diffusion Coefficients in Sieved Soils in Relation to Water Content and Bulk Density. Journal of Soil Science 1981, 32 (2), 225–236. https://doi.org/10.1111/j.1365-2389.1981.tb01702.x.

(34) Hashitani, T.; Tanaka, K. Measurements of Self-Diffusion Coefficients of the Nitrate Ion in Aqueous Solutions of Potassium Nitrate and Calcium Nitrate. J. Chem. Soc., Faraday Trans. 1 1983, 79 (8), 1765. https://doi.org/10.1039/f19837901765.

(35) Owen, A. G.; Jones, D. L. Competition for Amino Acids between Wheat Roots and Rhizosphere Microorganisms and the Role of Amino Acids in Plant N Acquisition. Soil Biology and Biochemistry 2001, 33 (4), 651–657. https://doi.org/10.1016/S0038-0717(00)00209-1.

(36) Kabala, C.; Karczewska, A.; Gałka, B.; Cuske, M.; Sowiński, J. Seasonal Dynamics of Nitrate and Ammonium Ion Concentrations in Soil Solutions Collected Using MacroRhizon Suction Cups. Environ Monit Assess 2017, 189 (7), 304. https://doi.org/10.1007/s10661-017-6022-3.

(37) Okano, Y.; Hristova, K. R.; Leutenegger, C. M.; Jackson, L. E.; Denison, R. F.; Gebreyesus, B.; Lebauer, D.; Scow, K. M. Application of Real-Time PCR To Study Effects of Ammonium on Population Size of Ammonia-Oxidizing Bacteria in Soil. Appl Environ Microbiol 2004, 70 (2), 1008–1016. https://doi.org/10.1128/AEM.70.2.1008-1016.2004.

(38) Prosser, J. I. Autotrophic Nitrification in Bacteria. In Advances in Microbial Physiology; Rose, A. H., Tempest, D. W., Eds.; Academic Press, 1990; Vol. 30, pp 125–181. https://doi.org/10.1016/S0065-2911(08)60112-5.

(39) Jiang, Q. Q.; Bakken, L. R. Comparison of Nitrosospira Strains Isolated from Terrestrial Environments. FEMS Microbiology Ecology 1999, 16.

(40) Belser, L. W. Population Ecology of Nitrifying Bacteria. Annu. Rev. Microbiol. 1979, 33 (1), 309–333. https://doi.org/10.1146/annurev.mi.33.100179.001521.

(41) Geets, J.; Boon, N.; Verstraete, W. Strategies of Aerobic Ammonia-Oxidizing Bacteria for Coping with Nutrient and Oxygen Fluctuations. FEMS Microbiology Ecology 2006, 58 (1), 1–13. https://doi.org/10.1111/j.1574-6941.2006.00170.x.

