## Supporting Information for "Benefits and limits of biological nitrification inhibitors for plant N uptake and the environment"

**Contents**

SI Text:
- Model description
- Model parameterization
- Simulation

SI Figures: S1 and S2
SI References

### Model description

We simulate spatio-temporal changes in concentrations of BNIs, ammonium, nitrate, and the population dynamics of nitrifiers in the rhizosphere. Roots can exude BNIs and take up ammonium and nitrate through their – by root hairs expanded – surface area. Nitrifiers use ammonium for both the construction of proteins (immobilization) and the conversion to nitrate (nitrification) (graphical abstract and Figure 2). The oxidation of ammonium to nitrite is assumed to be the rate-limiting process of nitrification, and nitrite accumulation is not considered. Therefore, we model nitrification as ammonium conversion to nitrate by a group of nitrifiers in the rhizosphere.

We define an average rhizosphere NUE for the rooted zone in a field as

$$\text{NUE} = \frac{\text{uptake } NH_4^+ \text{+uptake } NO_3^- \text{ (µmol)}}{\text{total initial inorganic rhizosphere N (µmol)}}, \quad (1)$$

We define a relative N loss to the environment (RNL) per initial amount of N in the rhizosphere

as

$$\text{RNL} = \frac{\text{lost } NO_3^- \text{ from rhizosphere (µmol)}}{\text{total initial inorganic rhizosphere N (µmol)}}, \quad (2)$$

which is an average net N loss from the rooted zone in a field.

#### Ammonium and nitrate transport-reaction equations

Ammonium concentration in soil solution, $A_\ell$, is the substrate that is enzymatically oxidized by nitrifying bacteria, which use the energy for life processes [1]. The change in ammonium concentration in soil can be described as the sum of changes caused by diffusion, advection, uptake by bacteria and plant root surfaces in the ecto-rhizosphere and rhizoplane, nitrification, and net ammonification:

$$b_A \frac{\partial A_\ell}{\partial t} = \underbrace{\frac{1}{r}\frac{\partial}{\partial r}\left(rD_A f\theta \frac{\partial A_\ell}{\partial r} + v_0 r_0 A_\ell\right)}_{\text{transport}} - \underbrace{I_A p(t)}_{\substack{\text{uptake}\\ \text{root hairs}}} - \underbrace{\frac{1}{Y}\gamma X}_{\substack{\text{uptake}\\ \text{nitrifiers}}} - \underbrace{qX}_{\substack{\text{nitri-}\\ \text{fication}}} + \underbrace{a}_{\substack{\text{ammoni-}\\ \text{fication}}}, \quad (3)$$

where $X$ is the nitrifier population density in the soil, the arrival time of the root ($t_R = 14$) is described by switching $p(t) = 0$ to 1 when $t \geq t_R$. The soil buffer power for ammonium is $b_A$, defined as constant $b_A = dA/dA_\ell$ and thus the ammonium concentration in soil is $A = b_A A_\ell$. The net ammonification rate is $a = 0$ for simplification. The nitrifier yield constant is $Y$, which represents an average C:N ratio over the functional group of nitrifiers because it relates the units of $X$ to $A_\ell$. The relative nitrification rate is

$$q(A_\ell) = q_{\max} \frac{A_\ell}{K_m + A_\ell} \cdot f_{\text{in}}(\text{BNI}_\ell), \quad (4)$$

where the saturation constant for oxidation is $K_m$, and maximum oxidation rate constant is $q_{\max}$. For the sake of parsimony, the modeled rates did not vary with temperature or water, and the N in microbial biomass did not become plant-available during the season. The nitrification inhibition function, $f_{\text{in}}$, is described below, and depends on the BNI concentration in soil solution, $\text{BNI}_\ell$. The intake of ammonium by root hairs is described by the sink-term $I_A(r)$, if present at distance $r$. Root hair uptake causes a rapidly forming diffusion profile, calculated as in [2]. BNIs may reduce the oxidation by a factor between 0 and 1, which is modeled as function, $f_{\mathrm{in}}$, depending on BNI concentration. The change in nitrate concentration in soil, $N$, is expressed in concentration in soil solution, $N_\ell$,

$$b_N \frac{\partial N_\ell}{\partial t} = \underbrace{\frac{1}{r}\frac{\partial}{\partial r}\left(r D_N f \theta \frac{\partial N_\ell}{\partial r} + v_0 r_0 N_\ell\right)}_{\text{transport}} - \underbrace{I_N p(t)}_{\substack{\text{uptake}\\ \text{root hairs}}} + \underbrace{qX}_{\substack{\text{nitri-}\\ \text{fication}}} + \underbrace{l N_\ell}_{\text{loss}}, \quad (5)$$

where $N = b_N N_\ell$. The sink terms are root hair uptake, $I_N$, and N loss to the environment, $lN_\ell$. Michaelis-Menten kinetics describe the uptake of ammonium and nitrate by the root for all types of transporters. For nitrate, the flux at the root surface is described by (inner-boundary condition at $r = r_0$)

$$D_N f \theta \frac{\partial N_\ell}{\partial r} + v_0 N_\ell = \frac{V_{max,N}(N_\ell - N_{min})}{K_{m,N} + N_\ell - N_{min}} p(t). \quad (6)$$

And the flux at the mid-distance to a mirrored neighboring root segment is (zero-flux outer-boundary condition at $r = r_1$)

$$D_N f \theta \frac{\partial N_\ell}{\partial r} + \frac{v_0 r_0}{r_1} N_\ell = 0\,. \quad (7)$$

The boundary equations for ammonium are equivalent. The initial concentrations in soil solution are constant, $A_{\ell,\mathrm{init}}$ and $N_{\ell,\mathrm{init}}$, at $t = 0$.

#### Biological nitrification inhibition by root exudates

In the nitrification pathway, the rate-limiting process of chemolithoautotrophs is mostly the first step by the enzyme ammonia monooxygenase [3,4]. Thus, if oxidation is described by a Monod equation, inhibition could be described in different ways: 1) using steady-state enzyme-inhibition kinetics for oxidases; or 2) empirically by scaling the limit $q_{max}$ or 3) by reducing the slope via $K_m$. Since studies on BNIs give values for the fractional reduction of nitrification the Monod equation (approach 2) was scaled with a fitted reverse saturation function (between one and zero). The function

$$f_{\mathrm{in}}(\mathrm{BNI}) = \frac{\mathrm{K_{in}}}{\mathrm{K_{in}} + \mathrm{BNI}^2} \quad (10)$$

was fitted against inhibition data from literature (Figure 1). The function $f_{\mathrm{in}}$ is unit-less and applied to $q$ and $\gamma$ to inhibit both growth and conversion to nitrate [4]. The root and root hairs are modeled to exude BNIs in the range of the function $f_{\mathrm{in}}$ (Figure 2e).

The transport equation in the rhizosphere and accompanying initial-boundary conditions for the BNI concentration are

$$b_{\mathrm{BNI}} \frac{\partial \mathrm{BNI}_\ell}{\partial t} = \frac{1}{r}\frac{\partial}{\partial r}\left(r D_{\mathrm{BNI}} f \theta \frac{\partial \mathrm{BNI}_\ell}{\partial r} + v_0 r_0 \mathrm{BNI}_\ell\right) + E_{\mathrm{h}}\, p(t) - l_{\mathrm{deg}} \mathrm{BNI} \quad (11)$$

$$D_{\mathrm{BNI}} f \theta \frac{\partial \mathrm{BNI}_\ell}{\partial r} + v_0 \mathrm{BNI}_\ell = F_{\mathrm{ex}}\, p(t) \qquad \text{at } r = r_0, t > 0 \quad (12)$$

$$D_{\mathrm{BNI}} f \theta \frac{\partial \mathrm{BNI}_\ell}{\partial r} + \frac{v_0 r_0}{r_1} \mathrm{BNI}_\ell = 0 \qquad \text{at } r = r_1, t > 0 \quad (13)$$

$$\mathrm{BNI}_\ell = 0 \qquad \text{at } t = 0\,, \tag{14}$$

where $p(t) = 1$ for $t \geq t_R$, else $p(t) = 0$, and $\mathrm{BNI} = b_{\mathrm{BNI}}\mathrm{BNI}_\ell$. As with uptake, the exudation starts at the arrival time of the root, $t_R = 14$. $F_{\mathrm{ex}}$ is the exudation rate at the root surface $r_0$. The source-term of BNI from root hairs $E_{\mathrm{h}}$ is described over the volumetric root hair surface area: $E_{\mathrm{h}}(r) = \mathrm{A_h}(r)F_{\mathrm{ex}}$, for $r$ not larger than the average root hair length and $\mathrm{A_h}(r)$ describes the root hair surface per volume depending on the distance $r$. Over time, microbial degradation of BNI is possible but set to zero (i.e., not simulated): $l_{\mathrm{deg}} = 0$. Other factors can inhibit oxidation, like too much or little oxygen [5]. In this study, we exclude these effects and assume that limited oxygen and carbon dioxide do not further inhibit nitrification. For example, similar models investigated the inhibition of denitrification by oxygen in wetlands [6].

##### Nitrifier population

The previous partial differential equations are discretized such that the rhizosphere domain is divided radially into compartments. The density of microorganisms is modeled as ordinary differential equation (ODE) system. An ODE for each spatial compartment ($i = 1, \ldots, n$) from the numerical discretization of the substrate, resulting in a spatial gradient of nitrifier cell density

$$\frac{\partial X_i}{\partial t} = (\gamma(A_{\ell,i}) - \delta)X_i\,, \tag{8}$$

where $X$ is a vector with entries $X_i = X(r_i, t)$ and represents the total cell density of nitrifiers in soil defined as $X := b_X X_\ell$, with soil buffer power of nitrifiers, $b_X$, and nitrifier density in soil solution, $X_\ell$. Growth ($\gamma$) and death ($\delta$) hold for $X$, that means nitrifiers attached on soil particles are active, but they depend on the substrate in the solution $A_\ell$. The relative growth rate of the microbial population is described by

$$\gamma(A_\ell) = \gamma_{\max}\frac{A_\ell}{K_s + A_\ell} \cdot f_{\mathrm{in}}(\mathrm{BNI}_\ell), \tag{9}$$

where BNIs reduce the growth by a factor $f_{\mathrm{in}}$, $\gamma_{\max}$ is the maximum growth rate and $K_s$ is a saturation constant for growth of nitrifiers. The microbial death rate coefficient is denoted by $\delta$, and usually first-order, therefore, set constant. Motility of nitrifiers was not important to NUE and RNL [7], so that equation 8 is modeled as ODE system, where populations are still spatially distinct.

#### Model parameterization

The parameter values are within ranges based on the literature and given in Table 1. Most BNI research seems to focus on Poaceae, especially crops like rice, maize, and sorghum. Our reference parameterization is thus strongly based on data from these species, particularly rice. We regard our parameterization, however, species indifferent, especially because our sensitivity analysis covers a large parameter space, in which the niches of many species and genotype might be represented.

##### Nitrification, cell density and growth of nitrifiers

Parameters on specific oxidation can differ in their unit: relating to cells, *amoA* genes, or gram carbon as microbial biomass, and some publications did not normalize for bacterial concentrations but give rates of nitrate production in soil. We used maximum oxidation rate constant based on data of cell densities of nitrifiers. We used default values approximately in the middle of reported minimum and maximum values for $q_{max}$, $K_m$, $\gamma_{max}$, $K_s$, and $Y$. The reference value of initial cell density of nitrifiers was calibrated to agree with the oxidation rate coefficients and ammonium concentrations.

#### Exudation of BNIs

Chemically BNIs are a mixture of exudates consisting of various compounds, some hydrophobic others hydrophilic. Synergistic behavior of those different BNIs is suggested but not yet confirmed [3,8]. Hence, BNIs can differ in their solubility in water and, therefore, the apparent diffusion rates. Hydrophobic BNIs may concentrate at the rhizoplane, sticking to the root hair tips or main surface, whereas hydrophilic BNIs may diffuse. For the reference simulation, we used a moderate diffusion rate coefficient of BNIs in liquid, $D_{BNI}$, which is between the self-diffusion of water (order 1e-5 cm $s^{-1}$) and very slow diffusion (1e-9 cm $s^{-1}$). Note for the effective diffusion of BNIs in the soil solution (Figure 2e), we excluded adsorption to the soil solid and set the BNI soil buffer power to $\theta$.

Often, rates for exudation and maximum uptake rate coefficients are given as g per g root dry weight (DW) per unit time. Since such rates are estimated over the whole root system, they need to be understood as root system averages and do not account for local responses. Nevertheless, we used an average gram per root surface area factor to convert data and account for root diameter. Average factors for gram DW per rice root volume are calculated from the data [2]. The average g (DW) $cm^{-2}$ values are 0.00027, 0.00022, and 0.00034 for day 7, 14, and 21 after emergence, respectively. The values corresponding to g (DW) per root volume are 0.045, 0.038, and 0.065 for the same time points. For the used root radius and root hair parameters, this is 0.076 g per root volume, which is a plausible value [9] as an average conversion factors that does not account for root structure (fine root vs. thick roots).

The exudation rate was set concerning 10.8 mg $g^{-1}$ root $d^{-1}$ [3,10] and converted to $cm^{-2}$. However, when comparing the BNI concentration near the root and the inhibition function $f_{in}$, this rate might be an upper limit. Therefore, we reduced this rate for the water-soluble exudate compound, e.g., MHPP, by one-third. The reported values of the hydrophobic BNI were over four orders of magnitude lower [11]. The mobility of BNIs, thereby, remains uncertain.

#### Root and root hairs

Root and root hair geometries are based on data from L-type lateral roots of rice [2]: a root hair number on the root surface per unit length of 700 $cm^{-1}$, a root hair radius of 0.0008 cm, and an average root hair length of 0.015 cm. However, the root radius can also represent a small lateral root of another species. With a slightly larger mid-distance to neighboring roots, $r_1$, and no other lateral roots branching into the rhizosphere of unit length, the root system has a representative root length density (1.85 cm $cm^{-3}$).

#### Root uptake kinetics

Uptake is modeled with Michaelis-Menten kinetics for the ammonium and nitrate concentration ranges, including minimum concentrations for uptake ($A_{min}, N_{min} > 0$, Table 1). Data of uptake kinetics of the nitrification inhibiting 6 weeks old rice seedling, Wuyunjing7 (WYJ7), without the minimum concentration for uptake is used [11]. A minimum concentration for uptake means in the model that the plant can lose nutrients if the concentration at the root surface drops below that minimum concentration. Since the maximum uptake rates were in units per gram root DW, we estimated the uptake kinetics with the same gram to root surface area conversion factor described above for the BNI exudation. Note that WYJ7 inhibits nitrification, and Wuyujing3 (WYJ3) does not; WYJ3 even promotes nitrification [11,12].

#### Simulation

The simulations were performed with and without inhibition by exudation of BNIs and the reference parameter values in Table 1. The simulation duration was 150 days, representing a typical life cycle of an annual plant, and the first 14 days were considered without root as an initial period of nitrification when the root is not established yet. Hence, until day 14, the nitrifiers can grow freely, and if enabled, BNI exudation starts on day 14 (Figure 2a). In the sensitivity analysis, the initial N concentration was held constant except for Figures 4a and b. That means for the change in the soil buffer power, the initial concentration in soil solution was changed according to

$$A_{\ell,\mathrm{init},c_b} = A_{\ell,\mathrm{init}}/(c_b \cdot b_A), \tag{15}$$

where $c_b$ expresses the relative change in $b_A$. The total initial N changed for changing initial ammonium concentrations.

The sensitivity of NUE and RNL to changes in parameter values was analyzed by varying the parameter values relative to their default over a 16-fold change, 4-times smaller and larger than the reference values.

Compared to literature values [13–16], our nitrification rates of the reference simulation are within the range but at the lower end, calculated from the slope of the first 30 simulated days. For example, Mohanty et al. [16] applied three doses of 10 mM $NH_4$-N to a soil with 225 mg $kg^{-1}$ available N, and reported a range of potential nitrification rates as produced 0.49—0.65 mM $NO_3^-$ $g^{-1}$ soil $d^{-1}$. In agricultural soils, nitrification rate-ranges of 1—5 and 1—34 mg N $kg^{-1}$ soil $d^{-1}$ can be found [14]. Soil conditions, sorption, and initial concentration, influence nitrification rates dramatically.

The model was solved by a backward differentiation formula with variable order from 1 to 5 and a spatial step size of $\Delta r$ = 2e-3 cm [17,18]. We used relative and absolute tolerances of 1e-3 and 1e-6, respectively, for the solute concentrations and cell densities of nitrifiers as well as the cumulative N uptake by the root and nitrifiers (Figure 3). The time stepping was adaptive and starting and maximum time steps were set to 1e-8 and 0.1 d, respectively.

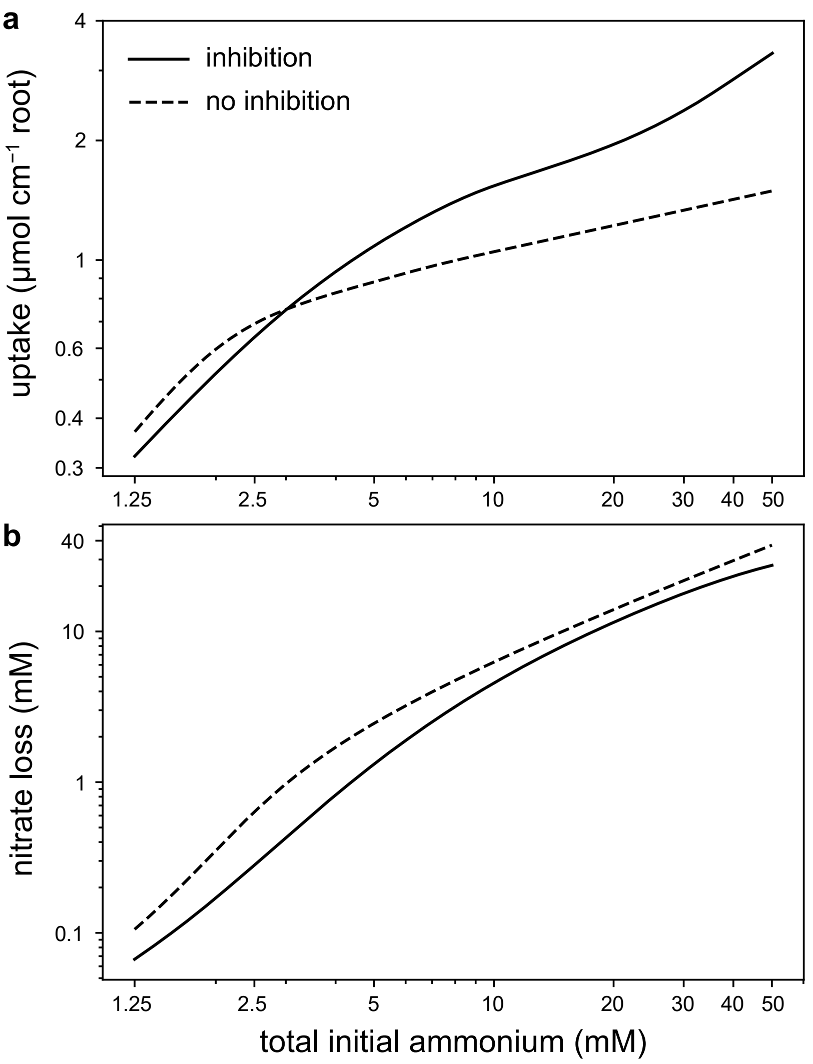


**Figure S1.** Influence of initial N in soil on (a) total N uptake per root segment of unit length and (b) nitrate concentration loss from the rhizosphere. Note, the axes are log-transformed.

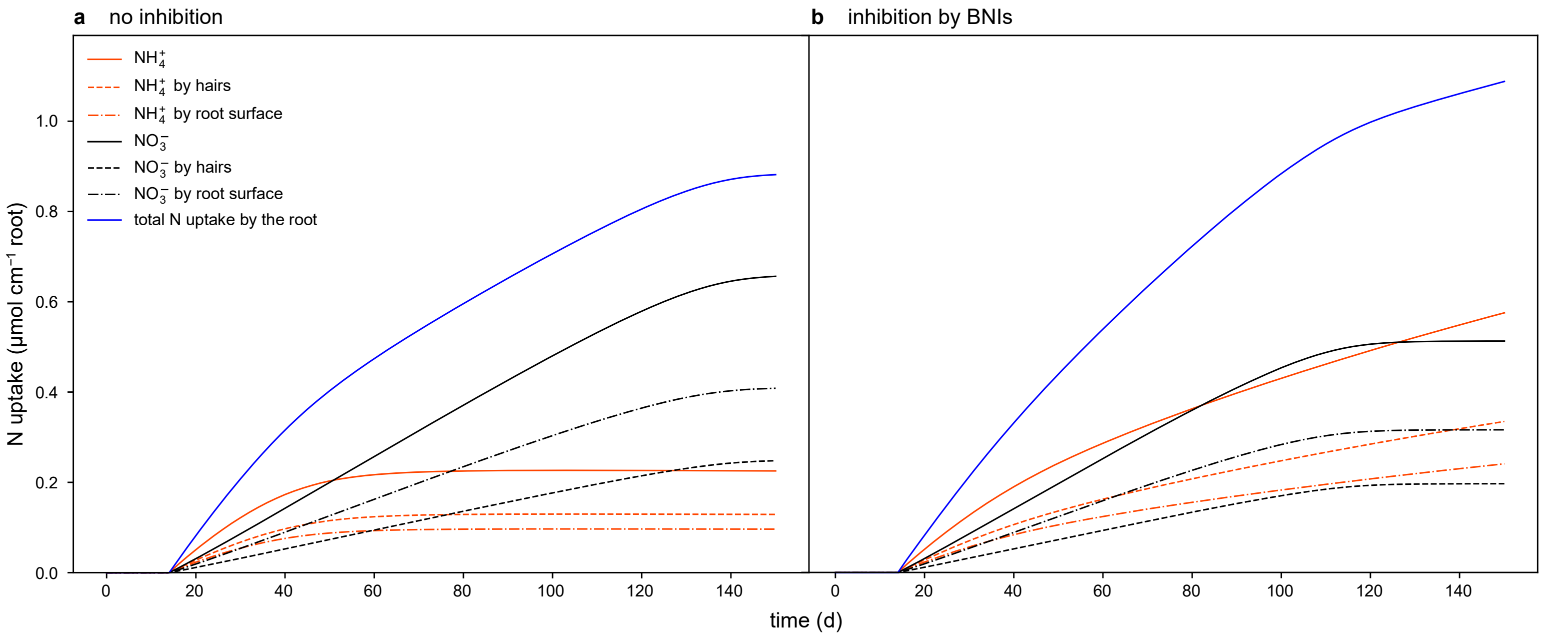


**Figure S2.** Cumulative uptake of nitrate and ammonium by a root of unit length and its hairs, without (a) and with BNIs (b).

#### SI References


(1) McLaren, A. D. Temporal and Vectorial Reactions of Nitrogen in Soil: A Review. *Can. J. Soil. Sci.* **1970**, *50* (2), 97–109. https://doi.org/10.4141/cjss70-017.

(2) Kuppe, C. W.; Kirk, G. J. D.; Wissuwa, M.; Postma, J. A. Rice Increases Phosphorus Uptake in Strongly Sorbing Soils by Intra-Root Facilitation. *Plant, Cell & Environment* **2022**, *45* (3), 884–899. https://doi.org/10.1111/pce.14285.

(3) Coskun, D.; Britto, D. T.; Shi, W.; Kronzucker, H. J. Nitrogen Transformations in Modern Agriculture and the Role of Biological Nitrification Inhibition. *Nature Plants* **2017**, *3* (6), nplants201774. https://doi.org/10.1038/nplants.2017.74.

(4) Nardi, P.; Laanbroek, H. J.; Nicol, G. W.; Renella, G.; Cardinale, M.; Pietramellara, G.; Weckwerth, W.; Trinchera, A.; Ghatak, A.; Nannipieri, P. Biological Nitrification Inhibition in the Rhizosphere: Determining Interactions and Impact on Microbially Mediated Processes and Potential Applications. *FEMS Microbiology Reviews* **2020**, *44* (6), 874–908. https://doi.org/10.1093/femsre/fuaa037.

(5) Prosser, J. I. Autotrophic Nitrification in Bacteria. In *Advances in Microbial Physiology*; Rose, A. H., Tempest, D. W., Eds.; Academic Press, 1990; Vol. 30, pp 125–181. https://doi.org/10.1016/S0065-2911(08)60112-5.

(6) Kirk, G. J. D.; Kronzucker, H. J. The Potential for Nitrification and Nitrate Uptake in the Rhizosphere of Wetland Plants: A Modelling Study. *Ann Bot* **2005**, *96* (4), 639–646. https://doi.org/10.1093/aob/mci216.

(7) Kuppe, C. W. Rhizosphere Models and Their Application to Resource Uptake Efficiency, RWTH Aachen University, Aachen, Germany, *unpublished*.

(8) Subbarao, G. V.; Nakahara, K.; Ishikawa, T.; Ono, H.; Yoshida, M.; Yoshihashi, T.; Zhu, Y.; Zakir, H. A. K. M.; Deshpande, S. P.; Hash, C. T.; Sahrawat, K. L. Biological Nitrification Inhibition (BNI) Activity in Sorghum and Its Characterization. *Plant Soil* **2013**, *366* (1), 243–259. https://doi.org/10.1007/s11104-012-1419-9.

(9) Shipley, B.; Vu, T.-T. Dry Matter Content as a Measure of Dry Matter Concentration in Plants and Their Parts. *New Phytologist* **2002**, *153* (2), 359–364. https://doi.org/10.1046/j.0028-646X.2001.00320.x.

(10) Zakir, H. A. K. M.; Subbarao, G. V.; Pearse, S. J.; Gopalakrishnan, S.; Ito, O.; Ishikawa, T.; Kawano, N.; Nakahara, K.; Yoshihashi, T.; Ono, H.; Yoshida, M. Detection, Isolation and Characterization of a Root-Exuded Compound, Methyl 3-(4-Hydroxyphenyl) Propionate, Responsible for Biological Nitrification Inhibition by Sorghum (Sorghum Bicolor). *New Phytologist* **2008**, *180* (2), 442–451. https://doi.org/10.1111/j.1469-8137.2008.02576.x.

(11) Sun, L.; Lu, Y.; Yu, F.; Kronzucker, H. J.; Shi, W. Biological Nitrification Inhibition by Rice Root Exudates and Its Relationship with Nitrogen-Use Efficiency. *New Phytologist* **2016**, *212* (3), 646–656. https://doi.org/10.1111/nph.14057.

(12) Chen, S.; He, M.; Zhao, C.; Wang, W.; Zhu, Q.; Dan, X.; He, X.; Meng, L.; Zhang, S.; Cai, Z.; Zhang, J.; Müller, C. Rice Genotype Affects Nitrification Inhibition in the Rhizosphere. *Plant Soil* **2022**. https://doi.org/10.1007/s11104-022-05609-9.

(13) Barber, S. A. *Soil Nutrient Bioavailability: A Mechanistic Approach*; John Wiley & Sons, 1995.

(14) Myrold, D. D.; Tiedje, J. M. Simultaneous Estimation of Several Nitrogen Cycle Rates Using 15N: Theory and Application. *Soil Biology and Biochemistry* **1986**, *18* (6), 559–568. https://doi.org/10.1016/0038-0717(86)90076-3.

(15) Højberg, O.; Binnerup, S. J.; Sørensen, J. Potential Rates of Ammonium Oxidation, Nitrite Oxidation, Nitrate Reduction and Denitrification in the Young Barley Rhizosphere. *Soil Biology and Biochemistry* **1996**, *28* (1), 47–54. https://doi.org/10.1016/0038-0717(95)00119-0.

(16) Mohanty, S. R.; Nagarjuna, M.; Parmar, R.; Ahirwar, U.; Patra, A.; Dubey, G.; Kollah, B. Nitrification Rates Are Affected by Biogenic Nitrate and Volatile Organic Compounds in Agricultural Soils. *Frontiers in Microbiology* **2019**, *10*.

(17) Shampine, L. F.; Reichelt, M. W. The MATLAB ODE Suite. *SIAM Journal on Scientific Computing* **1997**, *18* (1), 1–22. https://doi.org/10.1137/S1064827594276424.

(18) Kuppe, C. W.; Huber, G.; Postma, J. A. Comparison of Numerical Methods for Radial Solute Transport to Simulate Uptake by Plant Roots. *Rhizosphere* **2021**, *18*, 100352. https://doi.org/10.1016/j.rhisph.2021.100352.
